# Comparative circadian transcriptome analysis reveals dampened and phase-advanced rhythms in sun-exposed human skin

**DOI:** 10.1101/2025.09.30.679537

**Authors:** Michael M. Saint-Antoine, Zeyad El-Houni, Victoria L. Newton, Eleanor J. Bradley, Shruthi Ramesh, Hamish J.A. Hunter, Mike Bell, Alexander Eckersley, Michael J. Sherratt, Ron C. Anafi, Qing-Jun Meng

**Affiliations:** Department of Medicine, Chronobiology and Sleep Institute, Perelman School of Medicine, University of Pennsylvania, Philadelphia, PA 19104, US; Faculty of Biology Medicine and Health, The University of Manchester, Manchester Academic Health Science Centre, Manchester, UK; No7 Beauty Company, The Boots Group, Nottingham, UK; Medicines Evaluation Unit, Manchester, UK; Dermatology Centre, Northern Care Alliance NHS Foundation Trust, Manchester, UK; NIHR Manchester Biomedical Research Centre, Manchester University NHS Foundation Trust, Manchester Academic Health Science Centre, Manchester, UK

## Abstract

**Background:** Daily molecular rhythms modulate skin physiology. However, the effects of chronic sun exposure on these rhythms remain unstudied.

**Objectives:** This study aimed to identify and compare rhythmic genes and pathways in photoprotected and chronically photoexposed human skin *in vivo*.

**Methods:** Twenty healthy White women, aged 51-63, with moderate-severe photoageing of the dorsal forearm were recruited. Skin biopsies (3mm) were taken from photoprotected (upper buttock) and photoexposed (dorsal forearm) skin of each individual at noon, 6PM, midnight, and 6AM, across a 24-hour cycle. Skin biopsies were analysed by RNA sequencing. Cosinor analysis identified cycling genes along with their amplitudes and peak expression phases. Nested models were used to identify genes that were differentially rhythmic between the photoprotected and photoexposed sites. Phase set and gene set analyses identified pathways enriched among rhythmic transcripts or altered between the two sites.

**Results:** In the photoprotected buttock skin, 1546 genes (12%) met the criteria for cycling. In photoexposed forearm skin, the number was reduced to 959 (8%). As a group, transcripts that cycled in both sites had overall higher amplitude in photoprotected skin (*p* < 2.2e-16). Peak expression times for these transcripts showed a pronounced bimodal distribution and were clustered in the early morning and mid-afternoon. Distributions of peak times were significantly different between photoexposed and photoprotected skin (*p* < 0.00025), with peak times advanced in photoexposed skin. We identified 480 genes with significantly different rhythmic properties between the skin sites. Genes involved in DNA repair, MYC targets, E2F and G2M checkpoint pathways were enriched among those that showed higher amplitude oscillations in photoprotected skin. Genes involved in epithelial mesenchymal transition and apical junction pathways showed higher amplitude oscillations in photoexposed skin.

**Conclusions:** Temporal rhythms have a marked influence on skin molecular physiology and are altered in photoaged skin. Temporally advanced cycling patterns and a reduced number of rhythmic genes in photoexposed as compared to photoprotected skin suggest that chronic UV exposure may disrupt and/or reprogram circadian output rhythms to further alter skin physiology.

## Introduction

- One of the driving forces of the evolution of circadian clocks is to avoid irradiation while DNA is synthesised.
- Skin is one of the largest organs in the human body and interfaces with the rhythmic environment, such as daily exposure to temperature, humidity, and UV light.
- Many physiological parameters in skin experience diurnal changes, however, no studies have directly compared sun-exposed vs protected human skin *in vivo*.
- Understanding human skin chronobiology and how it adapts to and anticipates daily variations in stressors (e.g., UV light) is critical for the maintenance of skin health.

Many biological processes exhibit circadian (∼24 hourly) rhythms. In mammals, endogenous, self-sustained circadian clocks enable organisms to anticipate and adapt to diurnal changes associated with the light-dark cycle, including solar irradiation. These rhythms are organised hierarchically, with cell-autonomous oscillators found throughout the body. At the molecular level, a transcription-translation feedback loop (TTFL) drives the oscillatory expression of cell-specific targets, leading to rhythmic changes in physiology and behavior (1).

The skin, which is the largest organ in the human body, directly interfaces with the environment, including the rhythmic daily exposure to sunlight, pollutants, temperature, and humidity. As a result, skin can show distinct ageing phenotypes depending on its long-term exposure levels to extrinsic factors, most especially solar radiation, in particular UV radiation (UVR), which induces photoageing (2,3).

Intact skin tissue and isolated skin cells exhibit prominent diurnal rhythms. Zanello et al. demonstrated TTFL cycling in primary epidermal keratinocytes, dermal fibroblasts, and melanocytes (4). Numerous cellular processes show rhythms in intact skin, including keratinocyte proliferation, melanogenesis, dermal fibroblast migration, and epidermal stem cell differentiation (daily S-phase cycle) (5). Skin physiology, including immune regulation, temperature, water loss (TEWL) (6), and skin surface pH (7), as well as the hair follicle cycle (8), also exhibit prominent rhythms.

Transcriptomic analysis of primary keratinocytes has shown, as with other cell types and tissues, that different biological pathways peak at different times of the day. Pathways peaking in the early morning are proposed to promote differentiation, while those peaking in the evening are thought to serve as protective responses in anticipation of UV damage (9). Previous longitudinal studies in humans have collected skin samples at multiple timepoints and identified rhythmically expressed transcripts (10,11).

Despite these studies, it remains unclear how these rhythms are influenced by chronic sun exposure in human skin. Moreover, previous studies have focused on a younger demographic, including more males. This study aimed to: 1) identify rhythmic genes and pathways in human skin *in vivo* in older female volunteers, and 2) compare the diurnal transcriptional profiles of skin chronically exposed to solar radiation with those in skin predominantly protected from sun exposure.

## Materials and Methods

### Sample collection

We collected time course skin biopsies from twenty healthy White women, aged 51-63 (mean 57.5 years ±3.8) who exhibited moderate-severe photoageing of the dorsal forearm (Global McKenzie Scale Rating ≥ 4, **Table 1**). The study adhered to the guidelines of the International Council on Harmonization of Good Clinical Practice (12) and the recommendations of the Declaration of Helsinki (13). The study was approved by the Northwest – Preston Research Ethics Committee (ethics No: 10/H1016/25).

**Table 1:** Volunteer demographics. MSFsc – Mid Sleep on free days corrected for sleep debt accumulated during workdays. The McKenzie Scale Rating was performed on the entire dorsal forearm, fine wrinkles, coarse wrinkles and abnormal pigmentation were rated separately and used to generate a Global Assessment Severity Score (Mild: 1,2,3; Moderate:4,5,6; Severe: 7,8,9).

| Volunteer | Age | Chronotype | MSF<sub>SC</sub> | Dorsal Forearm Photoageing score<br>(Global McKenzie Scale Rating) |
| --- | --- | --- | --- | --- |
| 1 | 58 | More evening | 2.2 | 4 |
| 2 | 59 | More evening | 4.0 | 4 |
| 3 | 51 | Neither | 4.0 | 4 |
| 4 | 54 | More evening | 3.5 | 4 |
| 5 | 63 | More morning | 3.1 | 6 |
| 6 | 55 | Neither | 3.3 | 4 |
| 7 | 62 | More morning | 2.5 | 6 |
| 8 | 51 | Neither | 3.7 | 4 |
| 9 | 54 | Neither | 3.0 | 4 |
| 10 | 53 | Neither | 2.0 | 5 |
| 11 | 55 | Neither | 2.3 | 4 |
| 12 | 56 | Neither | 3.3 | 4 |
| 13 | 61 | Neither | 2.5 | 5 |
| 14 | 62 | More morning | 3.3 | 4 |
| 15 | 59 | More morning | 2.5 | 7 |
| 16 | 59 | More evening | 3.5 | 4 |
| 17 | 62 | Neither | 3.6 | 5 |
| 18 | 61 | More morning | 2.8 | 4 |
| 19 | 55 | More evening | 3.6 | 5 |
| 20 | 59 | More morning | 2.6 | 5 |

Biopsies were collected between 04Mar24 and 20Mar24, with volunteers at least one-year post-menopause, selected against the inclusion/exclusion criteria set out in **Table 2**. Volunteers were assessed for signs of wrinkling and pigmentation on the forearm as per the McKenzie Scale Rating and possessed either a moderate (4, 5, 6) or severe (7, 8, 9) global severity score as rated by a dermatologist. The buttock skin site was not formally scored, but none showed signs of photoageing.

**Table 2:**
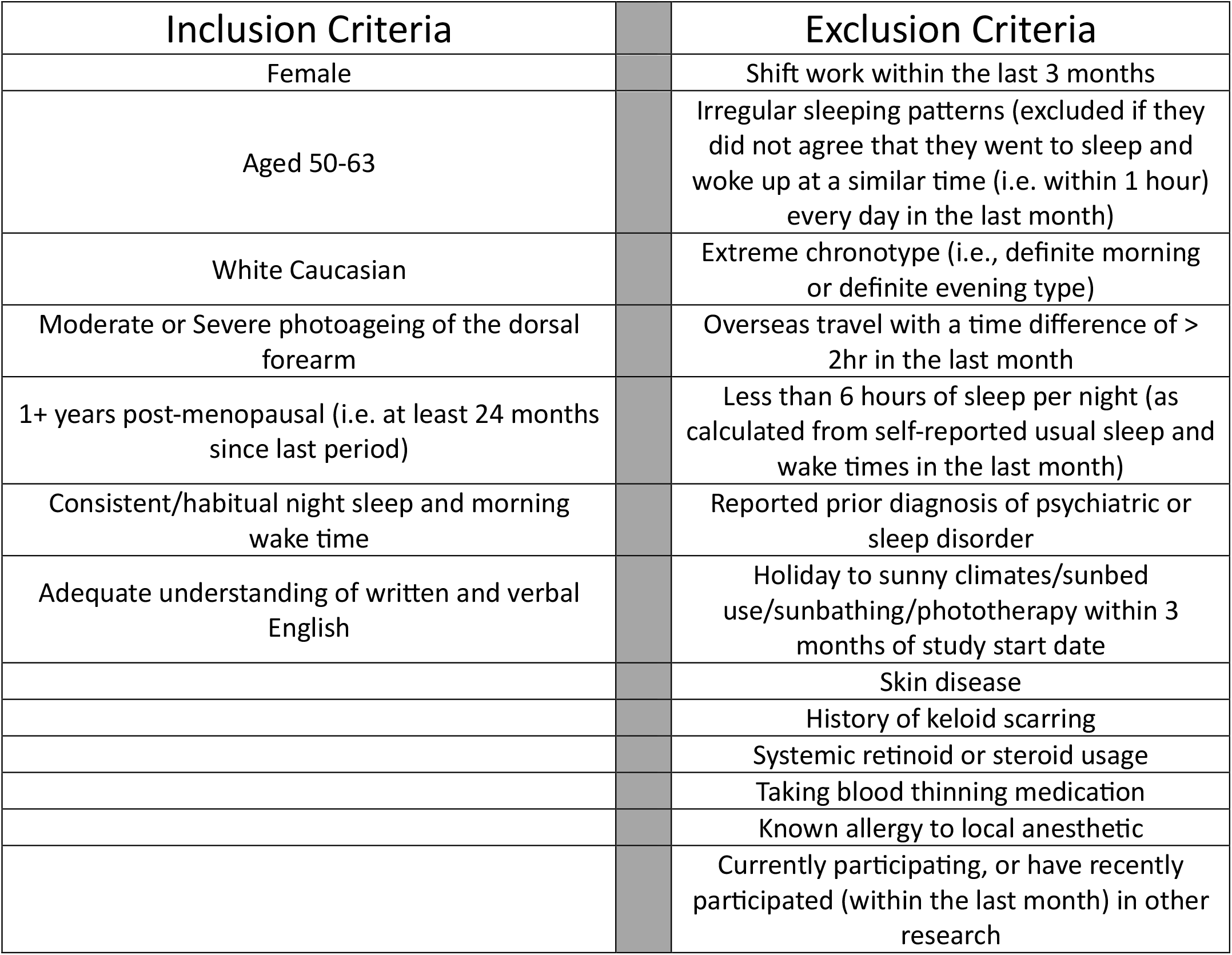
Study inclusion and exclusion criteria.

Volunteers were housed in a clinical facility for the duration of the study where they kept to their self-selected habitual sleep-wake times, excepting brief awakenings for biopsy collection. They were exposed to regular room lighting and were free to move around while they were awake. Full-thickness (3 mm) punch biopsies were collected every six hours during a single 24-hour period beginning at noon (*t = (12, 18, 0, 6))*. At each timepoint, two biopsies were obtained from each subject, one from the upper buttock (*PROTECTED*) and one from the dorsal forearm (*EXPOSED*). At each timepoint, buttock and forearm biopsies were obtained within 6 minutes of each other, with most (57/80) separated by less than a minute. Biopsies were immediately placed into RNAlater® and stored for 24-48 hours at 2-8°C prior to being frozen.

### Sequencing

Frozen biopsies were shipped on dry ice to Azenta Life Sciences (Genewiz, Germany) for RNA sequencing. RNA was extracted and purified using QIAsymphony RNA (Cat no. 931636; Qiagen, Hilden, Germany) and subjected to quality control prior to library preparation. 100ng of RNA per sample was used for library preparation, performed by polyA selection–based mRNA enrichment, mRNA fragmentation, and random priming with subsequent first- and second-strand complementary DNA (cDNA) synthesis. Following that, end-repair 5′ phosphorylation and adenine nucleotide (dA)–tailing was performed. Last, adaptor ligation, polymerase chain reaction (PCR) enrichment, and Illumina NovaSeq technology–based sequencing was performed with 20M reads per sample using Trimmomatic v.0.36 (14). Cleaned reads were aligned to the *Homo sapiens* GRCh38 reference genome using STAR aligner v2.5.2b with default parameters (15). This process yielded a raw counts dataset with measurements for 57,500 transcripts.

### Data processing

Statistical analysis was performed in R. Informatics packages were used with default parameters unless otherwise specified. RNA-seq raw counts were filtered to remove lowly expressed transcripts using the *filterByExpr()* function from *edgeR* (16). Batch effects from sample processing were corrected with ComBat-seq (17), specifying skin site (*PROTECTED* or *EXPOSED*) as a relevant biological variable. Data were then normalized with the *calcNormFactors()* function from *edgeR* and normalized counts were exported using the *cpm()* function (18). Transcripts with mean CPM < 5 across both conditions were excluded, leaving 12,471 transcripts for downstream analysis. Ensembl IDs were used as the primary transcript identifiers. IDs were mapped to official gene symbols using the *org*.*Hs*.*eg*.*db* R package (19) for convenience and gene set analysis. R scripts used for data processing and subsequent analyses are available at https://github.com/ranafi/exposed_vs_protected_skin.

### Identification of rhythmic transcripts

We used Cosinor regression (20,21) to identify rhythmic transcripts in each skin site separately. For each transcript in each condition, time-course transcript expression *x(t)* was fit to two nested linear models:

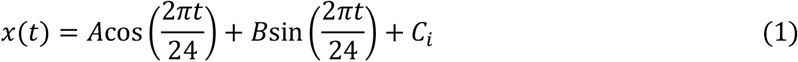

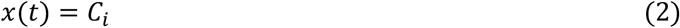

Here *i* ∈ {1,2, …,20} denotes one of the 20 subjects and *C*_*i*_represents the subject-specific circadian-corrected mean expression value or MESOR (midline estimating statistic of rhythm). Rhythmic parameters *A* and *B* were fit across all subjects collectively. The significance of rhythmicity was assessed with an F-test, using the *anova()* function, comparing the fit of the two models. P-values for each of the 12,471 transcripts were adjusted for multiple testing using the Benjamini-Hochberg false discovery method (22), and are referred to as q-values following adjustment.

The amplitude and acrophase (time of peak expression in hours) of each transcript were established using standard trigonometric identities:

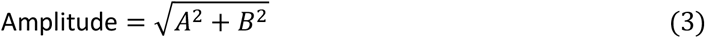

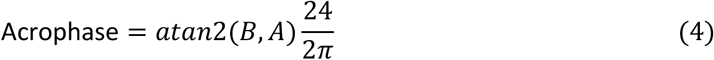

The relative amplitude for each transcript was calculated as the absolute expression amplitude divided by the mean of the subject-specific MESORs:

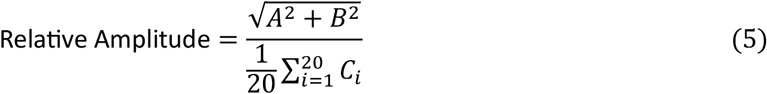

In the Results section, “amplitude” refers to relative amplitude. Our “first pass” analysis considered a transcript to be rhythmic if the Cosinor fit was significant (*q* ≤ 0.05) and the transcript had a relative amplitude ≥ 0.2.

### Differential rhythmicity testing

Transcripts identified as rhythmic in at least one site were tested for differential rhythmicity, again using a nested model approach. Combining data from both collection sites, we assessed the improvement in model fit when allowing separate rhythmic parameters for *EXPOSED* and *PROTECTED* skin as compared to simply allowing changes in mean transcript expression:

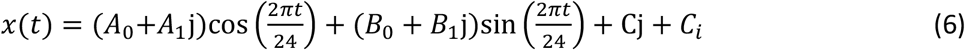

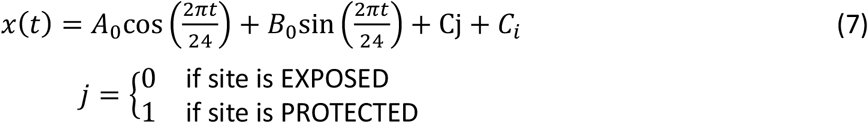

The improvement in model fit was again assessed using an F-test and results were corrected for multiple testing. Differentially rhythmic transcripts (*q* ≤ 0.1) were identified.

### Pathway analysis

Official gene symbols were used for set-level analysis. In cases where multiple Ensembl IDs mapped to the same symbol, the more rhythmic transcript (lower first-pass Cosinor p-value) was used. Gene sets from the Hallmark database (23) were used for initial pathway identification. Focused analysis of transcription factor targets was supplemented with lists from ENCODE (24) and ChEA (25).

We used Gene Set Enrichment Analysis (GSEA) (26) to identify pathways enriched with cycling transcripts. Genes were ranked by the negative log of their p-values from the first-pass Cosinor analysis. Ranked gene lists were analysed with the GSEA preranked tool. Phase Set Enrichment Analysis (PSEA) (27) identified pathways with temporally coordinated expression of rhythmic genes. Results from each skin site were analysed separately.

We also identified pathways that were enriched among transcripts that had higher or lower amplitude in each site. Genes identified as rhythmic in either skin site were ranked by the log ratio of their relative amplitudes in the two sites. The ranked list was analyzed with GSEA. To complement this analysis, we used the Enrichr web application (28) to conduct over-representation analysis. Genes identified as having statistically significant differential rhythmicity were analyzed, using all genes that were rhythmic in either site as the background set.

## Results

### 1,546 transcripts were identified as rhythmic in the PROTECTED skin site

Our first-pass Cosinor analysis identified 1,546 rhythmic transcripts in *PROTECTED* skin (**Table S1**) out of the 12,471 transcripts tested, including several canonical core clock genes, which we plotted for visual confirmation (**Fig. 1A**). As has been reported in other tissues (29,30), transcript acrophases were not uniformly distributed across the 24 hours. Rather, the distribution of transcript acrophases was bimodal, with two “rush hour” periods approximately 12 hours apart (**Fig. 1B**). PSEA analysis revealed that many pathways showed evidence of temporally coordinated expression (**Fig. 1C**) with the DNA repair pathway showing the greatest phase concentration.

**Figure 1:**
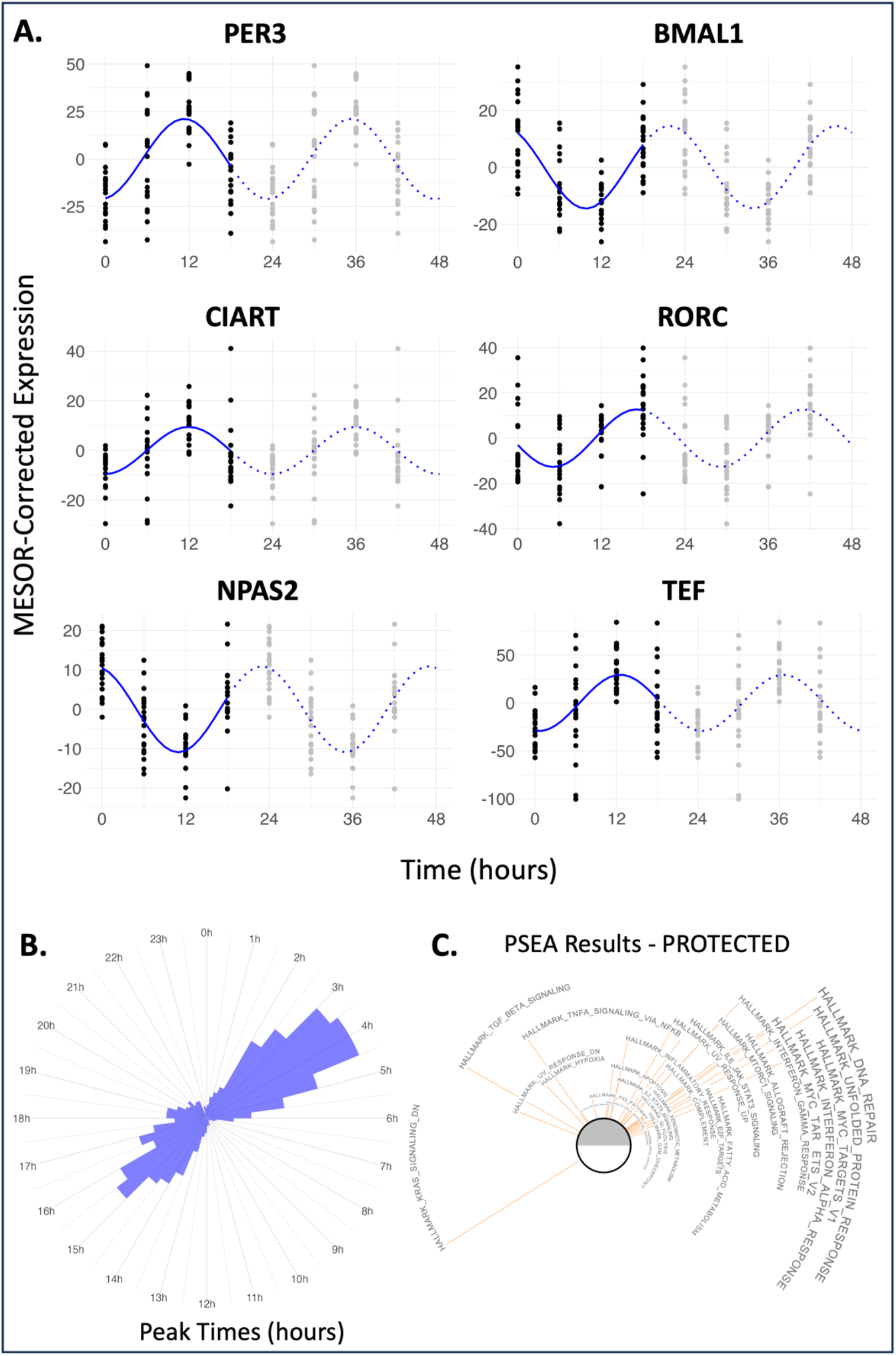
**A.)** MESOR-corrected timeseries expression plots of six core clock genes in the *PROTECTED* site, with fit Cosinor curves shown in blue (double-plotted for visualization purposes). **B.)** Acrophase (peak time) distribution for rhythmic genes in the *PROTECTED* site, plotted around a 24-hour clock. **C.)** PSEA results for the *PROTECTED* site. A similar plot for *EXPOSED* is available in **Fig. S1**.

### Fewer transcripts were identified as rhythmic in the EXPOSED skin site compared to the PROTECTED skin site

After analysing the *EXPOSED* skin sites in the same volunteers, we again visually confirmed the rhythmicity of select core clock genes (**Fig. 2A**). Although not every core clock gene met our pre-established criteria for rhythmicity (*q* ≤ 0.1 and relative amplitude ≥ 0.2) in both conditions, those that did showed closely aligned acrophases (**Fig. 2B**). The relative ordering of these transcripts also agreed with well-established molecular physiology from mouse models (31). Before considering differences between skin sites, we used a combination of GSEA set-level analyses to identify several pathways enriched for rhythmic genes in both sites (**Fig. 2C**), including MYC targets, unfolded protein response, and E2F targets.

**Figure 2:**
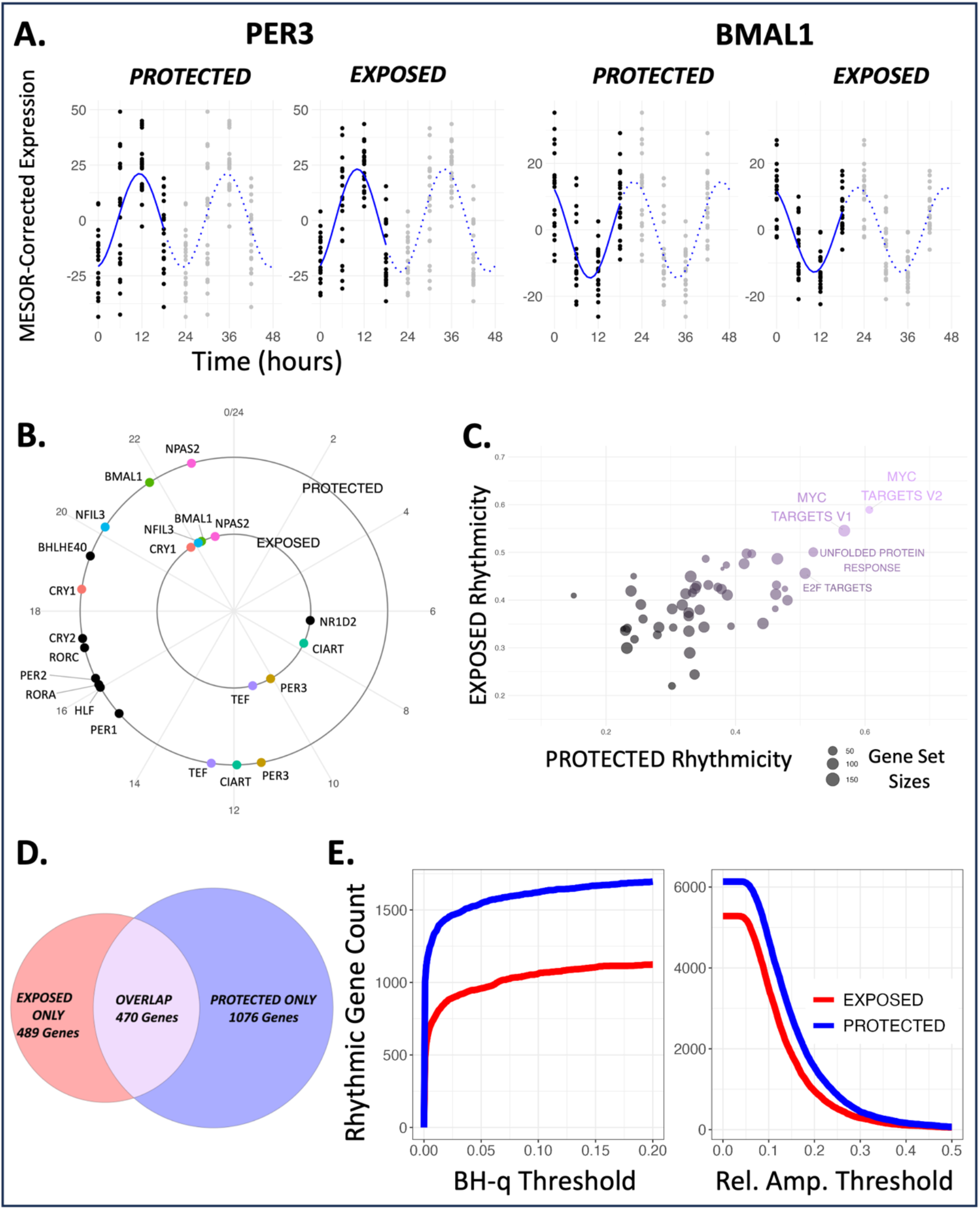
**A.)** MESOR-corrected timeseries expression plots for core clock genes PER3 and BMAL1, with fit Cosinor curves shown in blue (double-plotted for visualization purposes). **B.)** Core clock gene acrophase plots for *EXPOSED* (inner ring) and *PROTECTED* (outer ring). Genes that were rhythmic in both sites are color-matched. **C.)** GSEA enrichment scores for *EXPOSED* plotted against enrichment scores for *PROTECTED*. The pathways in the top right corner were found to be enriched for rhythmic genes in both conditions. **D.)** Venn diagram of rhythmic transcripts, for thresholds of *q* ≤ 0.05 and relative amplitude ≥ 0.2. **E.)** The pattern of fewer rhythmic transcripts in *EXPOSED* holds if we vary the q-value and relative amplitude thresholds.

Most strikingly, we noted that only 959 transcripts met our criteria for rhythmicity in *EXPOSED* skin (**Table S2**), a nearly 40% reduction as compared to *PROTECTED* skin (**Fig. 2D**). This pattern of fewer rhythmic transcripts in *EXPOSED* skin was robust to a range of different q-value and relative amplitude threshold choices (**Fig. 2E**).

### Rhythmic transcripts exhibited reduced amplitude and earlier peak time in the EXPOSED skin site compared to the PROTECTED skin site

We next focused on the 470 transcripts that were rhythmic in both skin sites (**Table S3**). For these genes, the distribution of amplitudes in *EXPOSED* skin was shifted leftward compared to the distribution in *PROTECTED* skin, suggesting that amplitudes were lower overall in *EXPOSED* skin (**Fig. 3A**). For each transcript, we computed the log-ratio of its amplitude in the two sites. The distribution of *log*_2_(*EXPOSED*/*PROTECTED*) values showed a leftward shift (one-sample *t*-test; mean = -0.167; *p* < 2.2e-16), again suggesting a pattern of reduced amplitude in *EXPOSED* skin (**Fig. 3B**). This pattern was also observed using absolute rather than relative amplitudes (**Fig. S1B,C)**, although the differences between sites were somewhat more modest (one-sample *t*-test; mean = -0.13; *p* = 4.722e-07).

**Figure 3:**
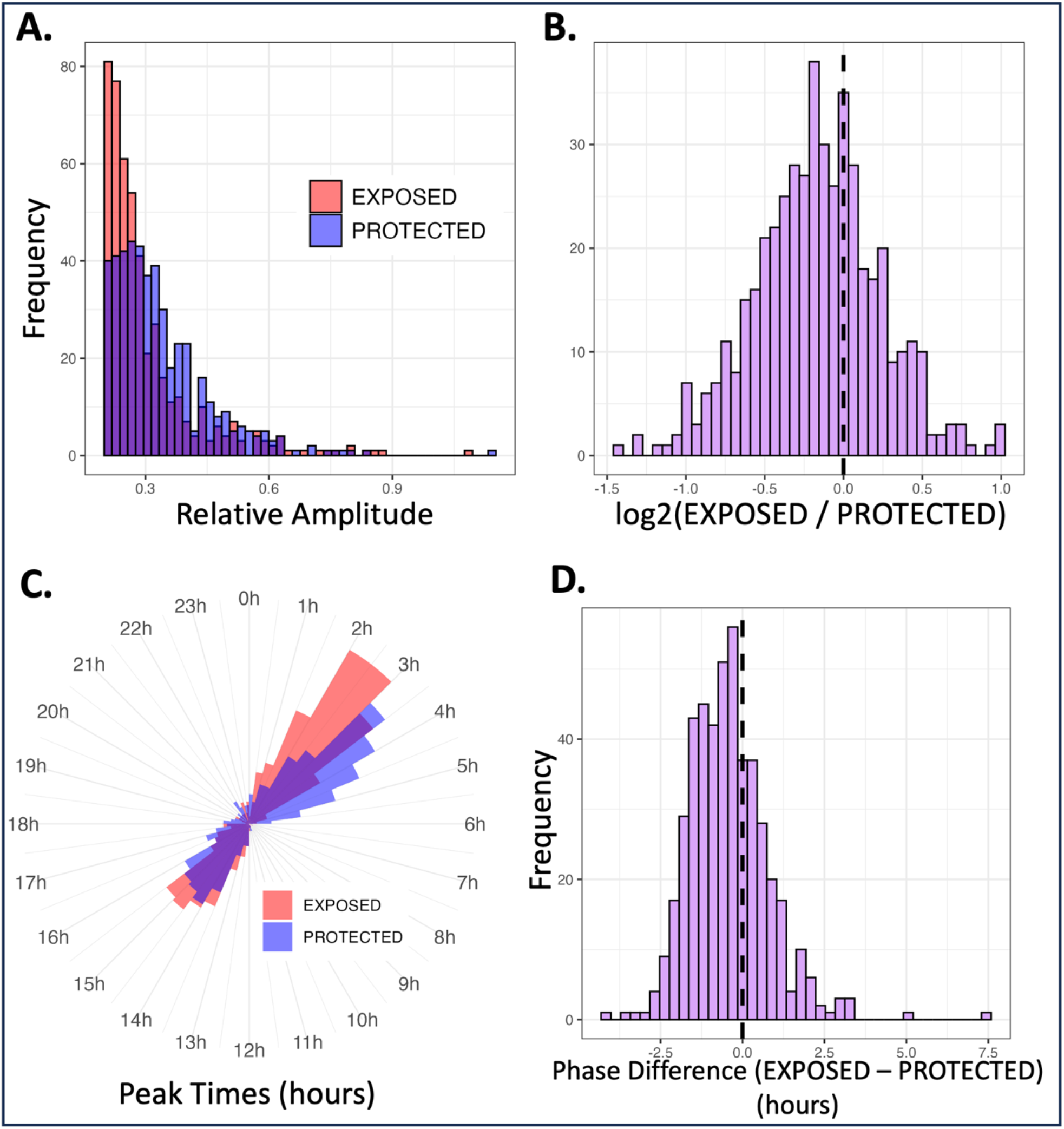
**A.)** Relative amplitude distributions in *EXPOSED* (red) and *PROTECTED* (blue), for the 470 transcripts that were rhythmic in both conditions. **B.)** Distribution of the *log*_2_-ratios of each transcript’s relative amplitude in *EXPOSED* to *PROTECTED*. Leftward shift indicates a pattern of lower amplitude in *EXPOSED* compared to *PROTECTED*. **C.)** Acrophase (peak time) distributions for *EXPOSED* and *PROTECTED* plotted around a 24-hour clock. **D.)** Distribution of the differences between acrophases (*EXPOSED* peak time minus *PROTECTED* peak time). Leftward shift indicates a pattern of earlier peak times in *EXPOSED* compared to *PROTECTED*.

In *EXPOSED* skin, almost two thirds (64%) of rhythmic transcripts peaked during only one third of the day, overnight between 10PM and 6AM. In *PROTECTED* skin, 51% of transcripts peaked between these hours (**Fig. S1D**). Considering again the set of 470 transcripts rhythmic in both sites, acrophase distributions differed significantly between the sites (two-sample Kuiper test; *p* < 0.00025), with peaks appearing earlier in the *EXPOSED* skin (**Fig. 3C**). The distribution of phase differences for each transcript (*EXPOSED* peak time minus *PROTECTED* peak time, **Fig. 3D**) had a circular mean of -0.43 hours (25th percentile: -1.34 hours, 75th percentile: 0.07 hours), suggesting that, on average, transcripts peaked 26 minutes earlier in *EXPOSED* skin (25th percentile: 1 hour 20 minutes earlier, 75th percentile: 4 minutes later).

### Many individual transcripts showed statistically significant differential rhythmicity between the skin sites

One approach for identifying transcripts with rhythmic differences between the sites would be to compare the lists of rhythmic transcripts in each site. However, this approach relies on small shifts in arbitrary thresholds leading to both false-positive and false-negative errors (e.g., a q-value shift from 0.049 to 0.051 when compared to an arrhythmic profile, or a relative amplitude difference of 0.21 versus 0.19 counting as a difference, while an amplitude shift from 0.2 to 0.5 goes undetected). Thus, recent studies recommend the use of differential rhythmicity tests that directly compare the rhythmic patterns between conditions (32–34).

Differential Cosinor analysis identified 480 transcripts with evidence of significant (*q* ≤ 0.1) differential rhythmicity between skin sites (24% of the 2,035 found to be rhythmic in either site) (**Table S4**). These rhythmic differences between sites were often visually striking, showing clear changes in amplitude (**Fig. 4A**) or acrophase (**Fig. 4B**). Heatmaps depicting temporal expression patterns for each of the 480 transcripts in both sites demonstrate blunted rhythms in *EXPOSED* skin (**Fig. 4C**). The general pattern of reduced amplitude in *EXPOSED* as compared to *PROTECTED* skin was even more pronounced among transcripts showing significant differential rhythmicity (**Fig. 4D**), and the pattern of earlier peaks in *EXPOSED* skin was again evident (**Fig. 4E**).

**Figure 4:**
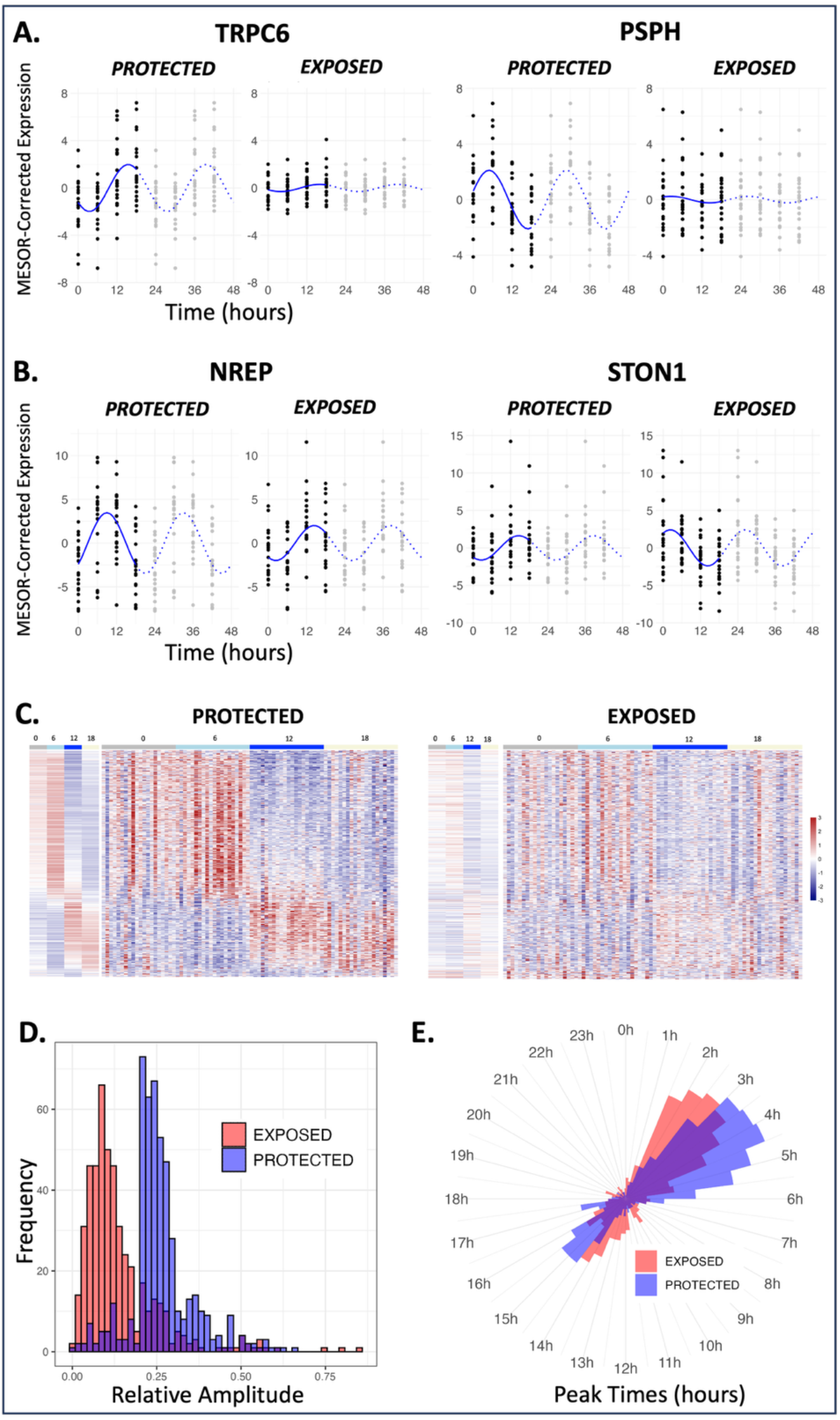
**A.)** MESOR-corrected timeseries expression plots for two differentially rhythmic genes that had large amplitude differences between the sites (double-plotted for visualization purposes). **B.)** MESOR-corrected timeseries expression plots for two differentially rhythmic genes that had large phase differences between the sites (double-plotted for visualization purposes). **C.)** Heatmap plots of z-score normalized expression for the set of 480 differentially rhythmic transcripts, in both *PROTECTED* and *EXPOSED*. Subplots to the left show timepoint means. **D.)** Relative amplitude distributions in *EXPOSED* (red) and *PROTECTED* (blue), for the 480 differentially rhythmic transcripts. **E.)** Acrophase distributions for *EXPOSED* and *PROTECTED* plotted around a 24-hour clock.

### The core molecular clock, DNA repair, and MYC pathways exhibited reduced amplitude in the EXPOSED skin site compared to the PROTECTED skin site

We next explored set-level differences in specific pathways: the core molecular clock, the DNA repair pathway, and the MYC pathway. All three have roles in proliferation, apoptosis, and cancer (35). DNA repair was the pathway that showed the largest magnitude of temporal concentration in *PROTECTED* skin (**Fig. 1C**), while the MYC pathway was among those identified by GSEA as being enriched for rhythmic genes in both sites (**Fig. 2C**). Lists of rhythmic genes in each of these pathways are in **Tables S5-7**.

We tested whether transcripts in these pathways, like the rhythmic transcriptome more generally, tended to have higher amplitudes in *PROTECTED* compared to *EXPOSED* skin. Indeed, this was true for all three pathways (two-tailed binomial; null proportion = 0.5; *p* < 0.05; **Fig. 5A**). Beyond simply having reduced amplitudes in *EXPOSED* skin, we then asked if this effect was stronger in these specific pathways as compared to the overall rhythmic transcriptome, in which 69% (1409/2035) of transcripts exhibited lower amplitude in *EXPOSED* compared to *PROTECTED* skin. MYC target genes stood out, showing a more marked trend for reduced amplitudes in *EXPOSED* skin than the rhythmic transcriptome at large (two-tailed binomial; null proportion = 0.69; *p* = 1.68e-15).

**Figure 5:**
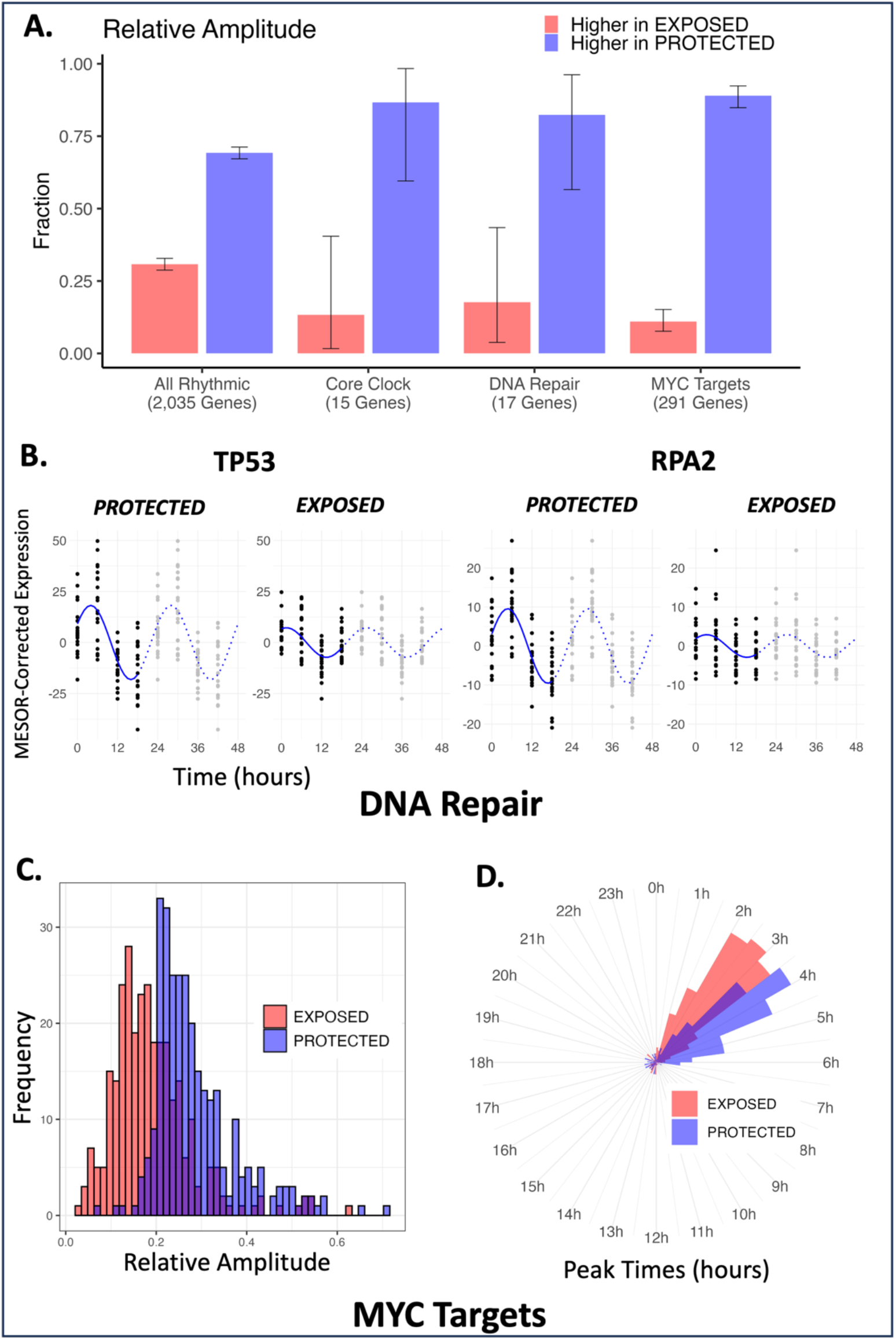
**A.)** Fractions of genes that had higher relative amplitude in each site, for the set of all rhythmic genes, rhythmic core clock genes, rhythmic DNA repair genes, and rhythmic MYC targets. Bars show Clopper–Pearson exact 95% confidence intervals. **B.)** MESOR-corrected timeseries expression of two DNA repair pathway genes (double-plotted for visualization purposes). **C.)** Relative amplitude distributions in *EXPOSED* (red) and *PROTECTED* (blue), for the 291 rhythmic MYC targets. **D.)** Acrophase distributions in *EXPOSED* and *PROTECTED* for the 291 rhythmic MYC targets.

As might be expected, despite these significant, aggregate pathway-level results, a more limited number of individual transcripts met significance criteria for differential rhythmicity. Of the 15 core clock genes rhythmic in either site, only CIART was significantly differentially rhythmic. Among the 17 oscillatory DNA repair genes, 6 were differentially rhythmic: TP53 (**Fig. 5B: left**), RPA2 (**Fig. 5B: right**), SAC3D1, IMPDH2, PDE4B, and NME1. Finally, 83 of the 291 annotated MYC targets were differentially rhythmic. The patterns of lower amplitude and earlier peaks in *EXPOSED* compared to *PROTECTED* skin are also clearly evident in MYC target expression (**Fig. 5C,D**).

### The G2M checkpoint and E2F pathways exhibited reduced amplitude in the EXPOSED skin site compared to the PROTECTED skin site

We used both enrichment and over-representation analysis to identify other pathways that had significantly higher amplitude in *PROTECTED* compared to *EXPOSED* skin (**Fig. 6A**). In addition to MYC targets, E2F targets (**Fig. 6B-D**) and G2M checkpoint (**Fig. 6E**) pathway transcripts showed a significant reduction in rhythmic amplitude in *EXPOSED* skin using both approaches. Lists of 462 rhythmic E2F targets and 43 rhythmic G2M pathway genes along with their circadian parameters are in **Tables S8**,**9**. E2F is a family of transcription factors involved in cell cycle regulation (36), DNA synthesis (36), and UV response (37).

**Figure 6:**
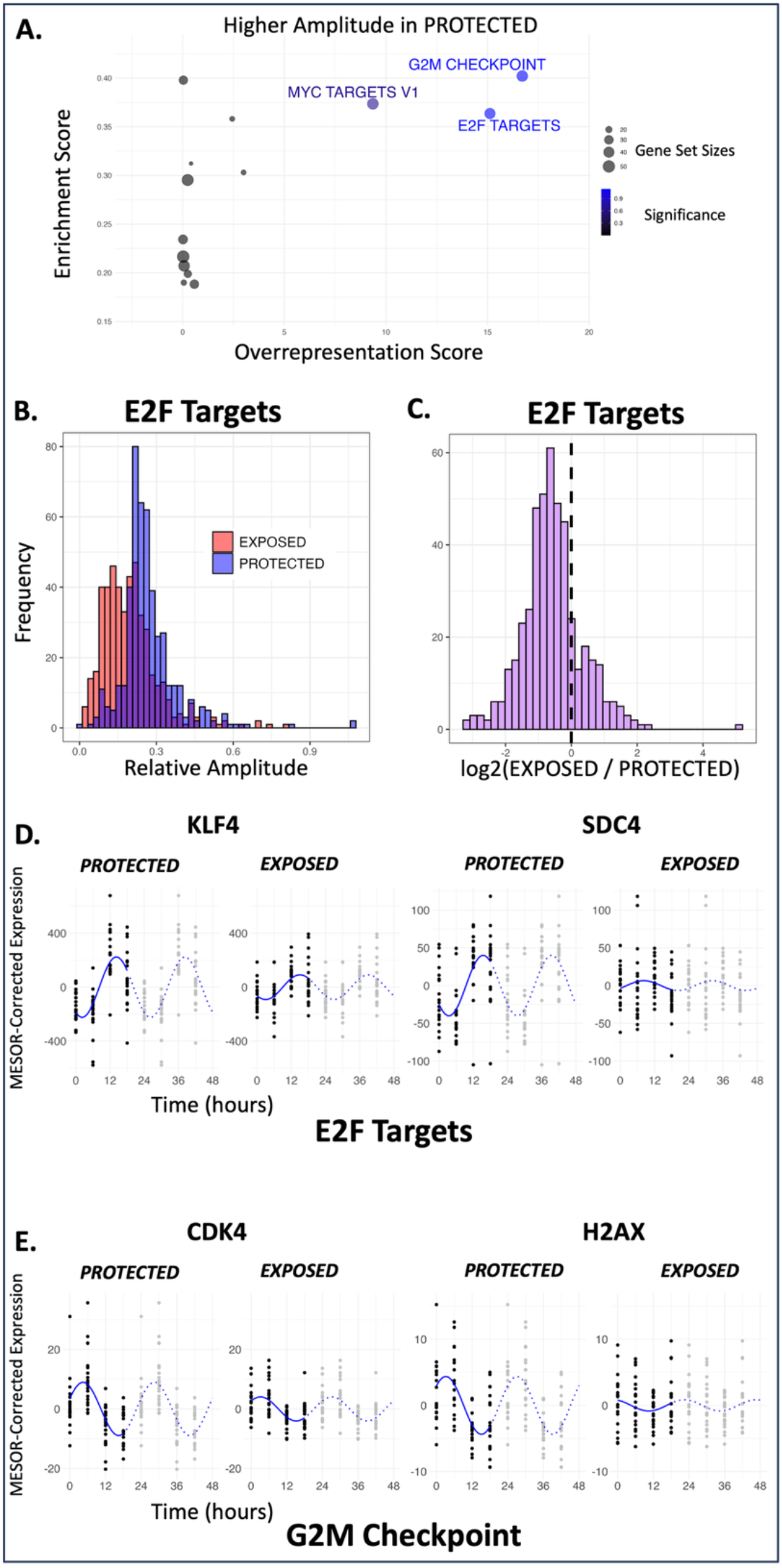
**A.)** Enrichment analysis (GSEA) results plotted against over-representation analysis (Enrichr) results. The top-right corner of the plot shows the consensus pathways that were identified by both methods as having higher overall amplitude in *PROTECTED* compared to *EXPOSED*. **B.)** Relative amplitude distributions in *EXPOSED* (red) and *PROTECTED* (blue), for the 462 rhythmic E2F targets. **C.)** Distribution of the *log*_2_-ratios of each transcript’s relative amplitude in *EXPOSED* to *PROTECTED*. **D.)** MESOR-corrected timeseries expression of two E2F targets (double-plotted for visualization purposes). **E.)** MESOR-corrected timeseries expression of two G2M checkpoint genes (double-plotted for visualization purposes).

Of the genes that constitute the E2F targets, KLF4, KLF9, CDC7, and SDC4 exhibited higher amplitude rhythms in the *PROTECTED* skin site compared to the *EXPOSED* skin site. The G2M/DNA damage checkpoint is a central component of cell cycle regulation that ensures cells do not enter mitosis (M-phase) with damaged DNA (38). In skin, this checkpoint plays a protective role by pausing the cell cycle to allow DNA repair in response to genotoxic stress such as ultraviolet radiation (39). Weaker rhythms in these pathways in *EXPOSED* skin suggest possible loss of circadian control of UV response, DNA synthesis, and cell cycle regulation due to chronic sun exposure.

### The epithelial mesenchymal transition and apical junction pathways exhibited increased amplitude in the EXPOSED skin site compared to the PROTECTED skin site

We next used the same combination of enrichment and over-representation approaches to identify pathways that had higher amplitude in *EXPOSED* compared to *PROTECTED* skin (**Fig. 7**). The epithelial mesenchymal transition (EMT) pathway (including IGFBP4, ABI3BP, PTX3, and THY1) and apical junction pathway (including VCAN, NLGN2, THY1, and ICAM1) were found to have increased amplitude in the *EXPOSED* skin site (**Fig. 7; Tables S10**,**11)**. EMT governs the loss of cell polarity and cell-cell adhesions along with an increase in migratory and invasive properties (40). Apical junctions are a hallmark of polarized multi-layered epithelial cells. They play a key role in maintaining the epidermal barrier, provide structural integrity, and act as a signaling hub within the skin (41). Our findings suggest that these pathways are reprogrammed in chronically UV-exposed, photodamaged skin, however, these correlations do not definitely establish UV exposure as the causal influence underlying altered rhythmicity.

**Figure 7:**
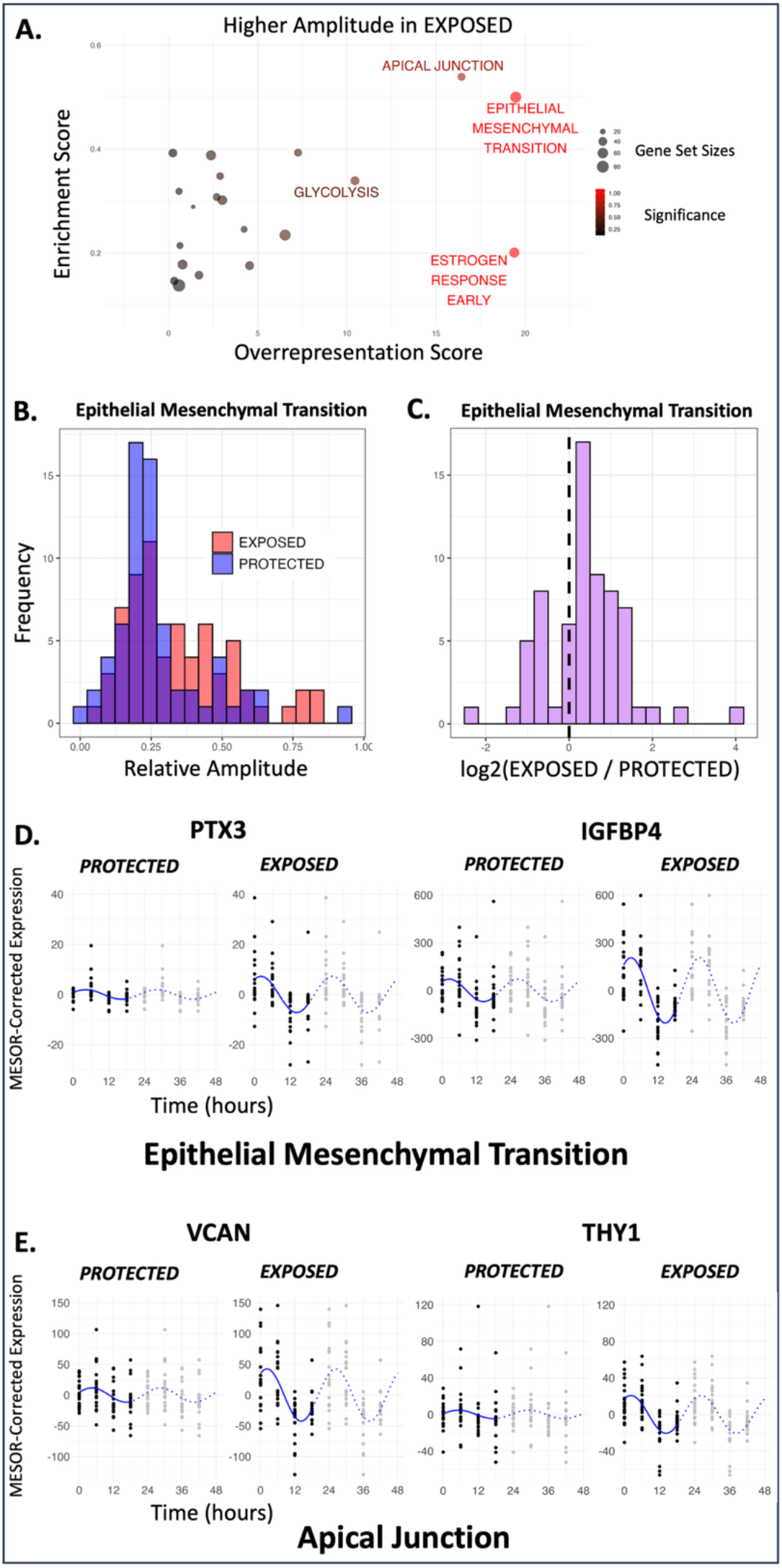
**A.)** Enrichment analysis (GSEA) results plotted against over-representation analysis (Enrichr) results. The top-right corner of the plot shows the consensus pathways that were identified by both methods as having higher overall amplitude in *EXPOSED* compared to *PROTECTED*. **B.)** Relative amplitude distributions in *EXPOSED* (red) and *PROTECTED* (blue), for the 67 rhythmic EMT-related genes. **C.)** Distribution of the *log*_2_-ratios of each transcript’s relative amplitude in *EXPOSED* to *PROTECTED*. Rightward shift indicates a pattern of higher amplitude in EXPOSED compared to PROTECTED **D.)** MESOR-corrected timeseries expression of two EMT-related genes (double-plotted for visualization purposes). **E.)** MESOR-corrected timeseries expression of two apical junction genes (double-plotted for visualization purposes).

## Discussion

Sun exposure is a key driver of extrinsic skin ageing (42). Exposure to excessive solar radiation, including UVR, causes acute (vasodilation driven erythema, swelling/inflammation, DNA damage, pain, etc.) and chronic (changes in skin pigmentation, mitochondrial dysfunction, alterations in the epigenome, changes to epidermal and dermal architecture, etc.) effects (43,44). Circadian clocks have a well-established role in skin physiology and skin clocks are being evaluated as a circadian biomarker (45,46). Using a statistically powerful within-subject design, this study identified and compared molecular rhythms in photoprotected and photoexposed human skin *in vivo*. We find circadian coordination of key pathways involved in the response to UV damage and daily stressors. Moreover, we find weaker clocks, phase-advanced rhythms, and fewer rhythmic genes in photoexposed skin, suggesting possible reprogramming of skin chronobiology following chronic UV exposure.

Nearly two thirds of transcripts cycling in sun exposed human skin peak at night, as compared to 51% in protected skin. Many pathways, including DNA repair, exhibit coordinated night-time peaks in transcript expression (**Fig. S1E**). UVR damages DNA, triggering a response that includes nucleotide-excision repair (NER) (47,48) and failure of the DNA repair pathway can lead to the development of cancer (49). The DNA repair pathway was previously identified as having a circadian rhythm in the mouse brain and liver (50), although the evolutionary role of this “nighttime repair” in humans remains an open question. Did circadian gating of DNA repair evolve as this process is more effective in the absence of UV exposure? Or does this gating anticipate and prepare the cell for coming molecular stressors? Indeed, with the delay between mRNA expression and protein translation, pathway protein activity might be well aligned with UV exposure.

We identified significant differences in circadian rhythmicity in photoexposed skin, including reduced amplitude and advanced phase. UVR-induced inflammation can result in degradation of fibrillin microfibril-rich oxytalan and elastic fibres and of fibrillar collagen, along with further remodeling of the dermal extracellular matrix (51–55). Mechanistically, in addition to direct cell-intrinsic effects (UV damage of DNA and micromolecules within cells), changes in ECM microenvironment of photoexposed skin may influence rhythmicity, leading to reduced amplitude, as seen in other epithelial tissues (56). Epigenetic changes may also be at play. In contrast to the overall trend, the EMT pathway (including matrix and remodelling genes such as Tenascin C, SERPINE1, and LOXL2) showed increased amplitude in photoexposed skin. Further studies are needed to investigate whether this reflects a functional change that provides a more effective response to UV-stress, or maladaptive UV-induced damage that favors more invasive and migratory cellular phenotypes.

Dorsal forearm and buttock skin biopsies have been frequently used in comparative studies of chronic UV exposure. Histological, biomechanical, and gene expression evidence support sun-exposure as the most important difference between these sites (57,58). Nonetheless, we cannot exclude the possibility that other factors contribute to the changes we observe. Similarly, we cannot distinguish oscillations directly driven by the core clock from those driven in response to daily rhythms in environmental exposures, food intake, or sleep/wake.

This study adds significant biological insight into the rhythmic genes and pathways in human skin. Through paired transcriptomic analysis, this work highlights the altered rhythms found in skin subjected to half a century of chronic UV exposure and is perhaps an additional feature of photoageing that one should strive to protect against with the use of sun protection measures. Our study has also revealed potential reprogramming of the genes and processes under circadian control in photoaged skin, which may allow it to better accommodate regular sun exposure.

## Acknowledgements

We thank No7 Beauty Company, part of The Boots Group, for funding this work. Thank you to the study participants for taking part in the study and the Medicines Evaluation Unit (MEU) study team for recruiting and obtaining the samples evaluated in this study. We would like to acknowledge Azenta Life Sciences (Genewiz) for carrying out the RNA extraction and sequencing of the samples. The views expressed are those of the authors and not necessarily those of the NIHR or the Department of Health and Social Care. The authors would like to thank Leo Zeef of the Bioinformatics and Genomic Technologies Core Facilities at the University of Manchester for providing support with the assessment of the quality of the RNA Sequencing data.

## Data Availability

The R scripts used for data processing and analysis are available at https://github.com/ranafi/exposed_vs_protected_skin. The transcriptomics data will be made available at ArrayExpress prior to publication.

## Ethics Statement

The study adhered to the guidelines of the International Council on Harmonization of Good Clinical Practice and the recommendations of the Declaration of Helsinki. The study was approved by the Northwest – Preston Research Ethics Committee (ethics No: 10/H1016/25).

---

**Figure S1:**
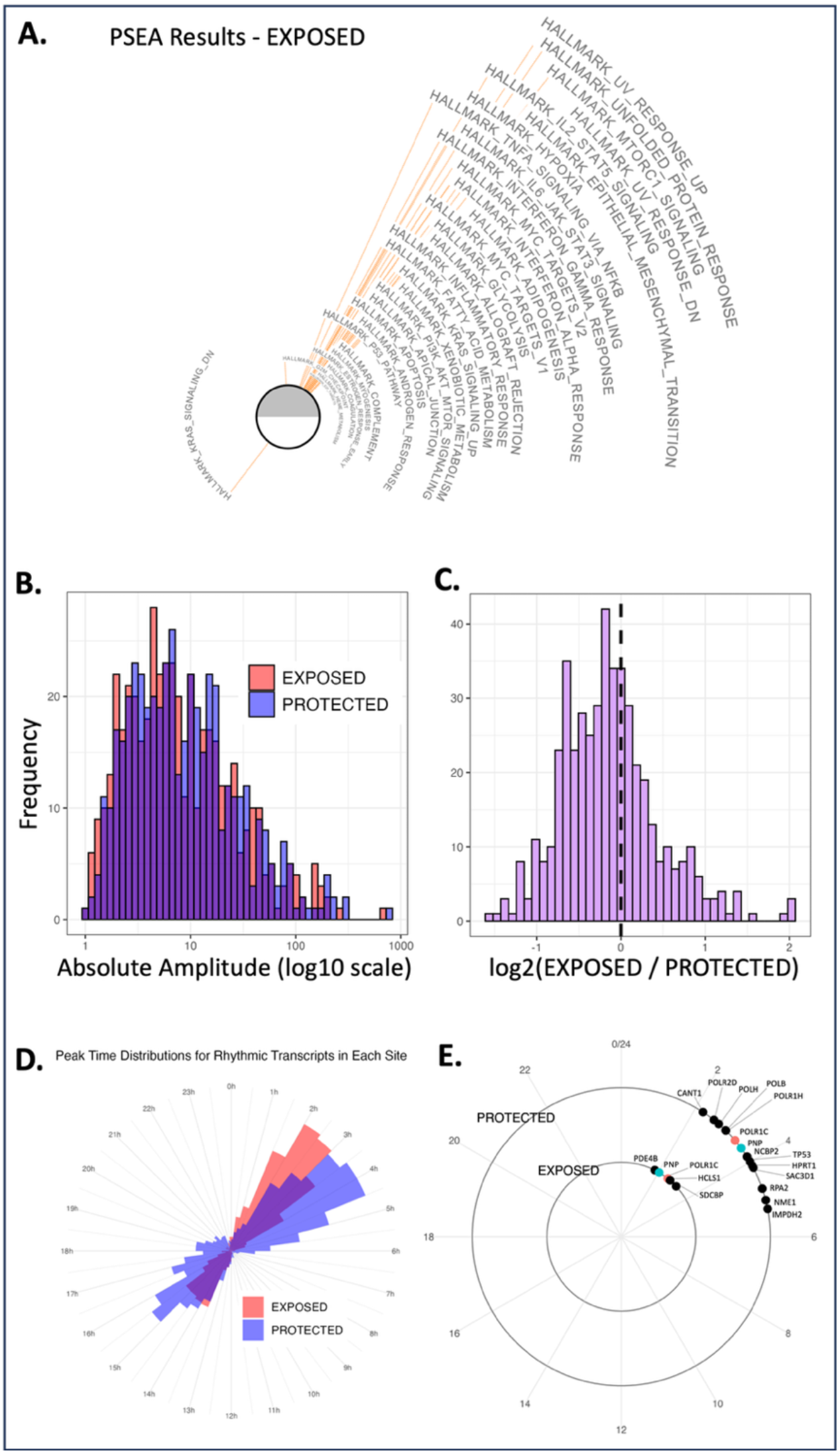
**A.)** PSEA results for the *EXPOSED* site. **B.)** Absolute amplitude distributions in *EXPOSED* (red) and *PROTECTED* (blue) for the 470 transcripts that were rhythmic in both conditions, with *log*_10_-scale horizontal axis, for visualization purposes. **C.)** Distribution of the *log*_2_-ratios of each transcript’s absolute amplitude in *EXPOSED* to *PROTECTED*. Leftward shift indicates a pattern of lower amplitude in *EXPOSED* compared to *PROTECTED*. **D.)** Acrophase (peak time) distributions for the set of rhythmic transcripts in each site. **E.)** DNA repair gene acrophase plots for *EXPOSED* (inner ring) and *PROTECTED* (outer ring). Genes that were rhythmic in both sites are color-matched. We note that more genes met the thresholds for rhythmicity in *PROTECTED* compared to *EXPOSED*, and that genes in both sites showed coordinated peak times in the early morning hours.

